# Auditory attention detection with EEG channel attention

**DOI:** 10.1101/2021.04.13.439732

**Authors:** Enze Su, Siqi Cai, Peiwen Li, Longhan Xie, Haizhou Li

## Abstract

Auditory attention detection (AAD) seeks to detect the attended speech from EEG signals in a multi-talker scenario, i.e. cocktail party. As the EEG channels reflect the activities of different brain areas, a task-oriented channel selection technique improves the performance of brain-computer interface applications. In this study, we propose a soft channel attention mechanism, instead of hard channel selection, that derives an EEG channel mask by optimizing the auditory attention detection task. The neural AAD system consists of a neural channel attention mechanism and a convolutional neural network (CNN) classifier. We evaluate the proposed framework on a publicly available database. We achieve 88.3% and 77.2% for 2-second and 0.1-second decision windows with 64-channel EEG; and 86.1% and 83.9% for 2-second decision windows with 32-channel and 16-channel EEG, respectively. The proposed framework outperforms other competitive models by a large margin in all test cases.

## I. INTRODUCTION

Humans have the ability to distinguish between speakers and to pay selective attention to one speaker in a multi-talker scenario, i.e., cocktail party [1]. However, hearing aid users experience the difficulty of following a target speaker in the presence of noise and other competing speech sources [2]. Are we able to equip the hearing aids with the human ability of selective attention? Recently, neuro-steered hearing prostheses are studied to produce a better experience for people with hearing loss, in which auditory attention is decoded from recordings of brain activity and used to enhance the speech separation for the attended speaker.

Among the brain signals for auditory attention detection (ADD), such as electrocorticographic (ECoG) [3], magnetoencephalography (MEG) [4] and electroencephalogram (EEG) [5], EEG is a more realistic option for brain-computer interface (BCI) applications, because it’s cheaper, non-invasive and easier to use. The techniques for EEG-based auditory attention detection can be grouped into linear and non-linear decoders [6].

Linear decoders have been well studied [5], where EEG responses are used to approximate the envelope of the speech heard by the participant, that is then compared with the original speech stimulus to reveal the attended or unattended speaker in a cocktail party scenario. Specifically, the reconstructed speech envelope from the cortical responses to a mixture of speakers is dominated by the salient spectral and temporal features of the attended speaker [3]. However, the correlation between the reconstructed and the attended envelope is fairly low [7]. A possible explanation is that the human auditory system is inherently non-linear [8] and the linear approach is probably not the best way to model the complex and dynamic nature of the brain [9]. Furthermore, the speech envelope reconstruction algorithm is not systematically optimized, e.g., jointly trained with the classifier, for auditory attention detection.

Recently, non-linear decoders have been studied to understand the complex and highly non-linear nature of auditory processes in human brain, that show superior performance to linear decoders [6], [7], [10], [11], [12]. In this paper, we follow the CNN-based non-linear approach [10], [11], [12] with a particular focus on low-latency settings. We note that the placement positions of electrodes reflect the activities of the related brain areas. Furthermore, some EEG channels are more informative than others in terms of informing the decision making process in the brain [13], [14]. At the same time, the distribution of effective channels may vary from subject to subject.

We propose a channel attention mechanism that predict a channel mask on-the-fly. The channel mask corresponds to a spatial map of the EEG electrodes, that gives a differentiated weight to each of the EEG channels. An element in the mask is a continuous value, as opposed to on-off channel selection, that modulates the contribution of each individual EEG channel for optimal auditory attention performance. Such a channel mask may vary with the attended speaker, the speech content, the acoustic environment, the listening subjects and so on. The question is how to devise a mechanism that dynamically predicts the mask according to each speech-EEG pair.

As far as we know, this is the first study on a channel attention mechanism for EEG-based auditory attention detection. The rest of the paper is organized as follows. Section II presents the channel attention mechanism and the CNN classifier. Experimental setup and the results are summarized in Section III. Finally, Section IV concludes the study.

## II. Auditory attention detection with channel attention

We study a CNN classifier with channel mask (CM) for AAD, which is referred to as CNN-CM hereafter, as illustrated in Fig. 1. The CNN-CM neural architecture consists of a channel attention mechanism and a CNN classifier.

**Fig. 1.**
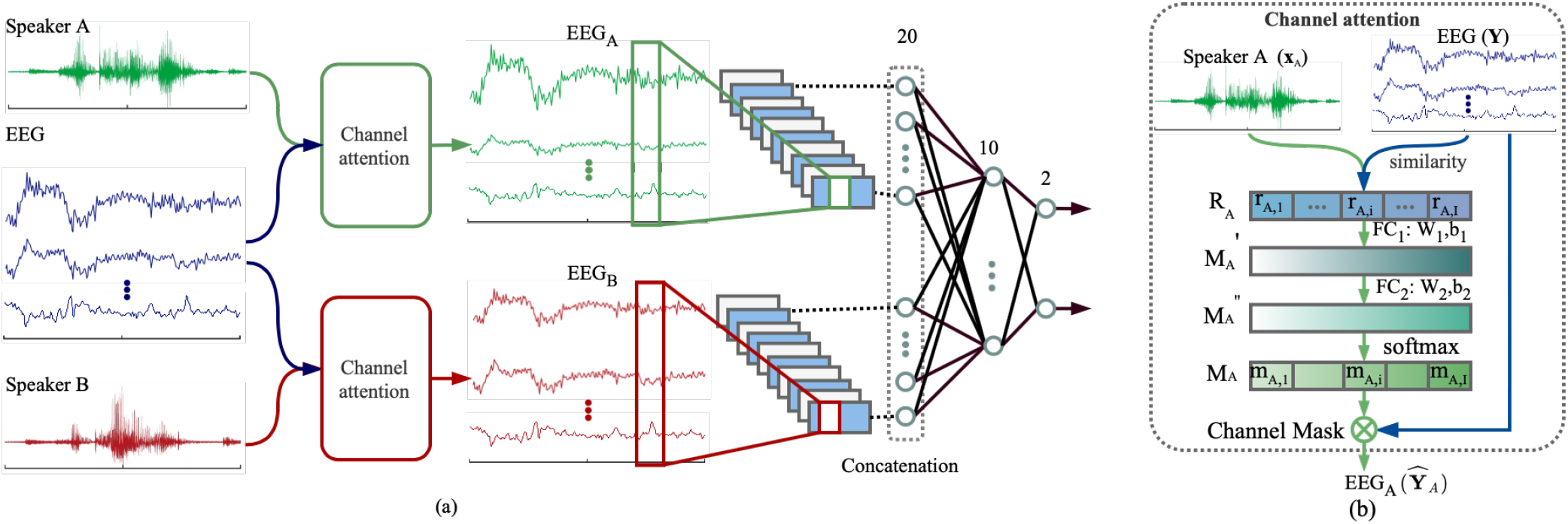
The proposed CNN-CM neural architecture for auditory attention detection, which is trained as a whole with two output nodes for two attended speakers. (a) Overall architecture; (b) Channel attention mechanism that modulates the input multi-channel EEG signals with respect to speaker *A*.

During training, the CNN-CM network takes multi-channel EEG signals, speech envelopes *A* and *B* as input, and the attention labels as the supervisory signals. The channel attention mechanism is trained to generate the modulated EEG signals via an attention mask, while the CNN classifier is trained to make a detection decision. Both the channel attention mechanism and the CNN classifier are jointly trained for optimal attention decision.

### A. EEG channel attention

Humans pay selective attention in many everyday situations, such as auditory attention in cocktail party scenarios. The channel attention mechanism is motivated by such human ability, that seeks to adaptively select important features in machine translation [15], image classification [16], [17] and caption generation [18]. In this study, we would like to dynamically assign weights to channels to reflect the contributions of individual EEG channels for AAD. The channel attention mainly has two properties, 1) it explicitly models the correlation between EEG responses and speech stimuli, and 2) adaptively adjusts the weights of EEG channels.

#### 1) Feature representation

The cosine similarity is chosen to measure the relationship between the speech stimuli and EEG responses [19] in this study, which does not involve any learning parameters. The cosine similarity between two time series, 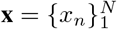 and 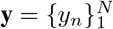, is defined as,

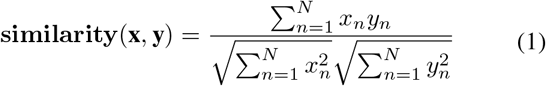

Let **x**_*A*_ be the speech envelope, i.e., the auditory stimulus, from speaker *A*. The correlation between **x**_*A*_ and the *i*^*th*^ channel of EEG signals, **y**_*i*_, can be denoted as,

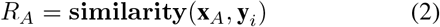

where *R*_*A*_ = [*r*_*A*,1_,*…, r*_*A,i*_,*…, r*_*A,I*_] and *r*_*A,i*_ denotes the correlation between speaker *A* and *I*-channel EEG signals. Similarly we can have *R*_*B*_ for speaker *B*.

#### 2) Predicting channel mask

In the neural attention mechanism, both *R*_*A*_ and *R*_*B*_ are taken as input by a gating mechanism to produce the channel mask {*M*_*s*_ : *s* ∈ {*A, B*}}.Two fully-connected (FC) layers are adopted to parameterize the gating mechanism to capture the nonlinear interaction among the channels [16]. The resulting channel mask for *I*-channel EEG signals 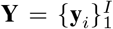 can be denoted as *M*_*s*_ =[*m*_*s*,1_, …, *m*_*s,i*_, …, *m*_*s,I*_],

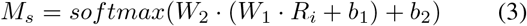

where a dimensionality-reduction layer with parameter *W*_1_ and bias *b*_1_ with reduction ratio *r* and *tanh* function as the activation function, and a dimensionality increasing layer with parameter *W*_2_ and bias *b*_2_, and followed by a sigmoid activation. Finally, the neural attention mechanism modulates the input EEG signals by applying the attention mask channel by channel 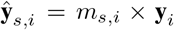 or 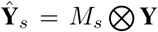 where ⊗ denotes a point-wise multiplication.

### B. Auditory attention detection

Convolutional neural network (CNN) has been employed as a classifier for auditory attention detection with state-of-the-art performance [6], [7], [10], [11], [12]. Hence, we adopt the CNN as a backend classifier that takes two sets of modulated EEG signals as the input, one for speaker *A* and another for speaker *B*, and decides which speech stimulus is associated with the EEG responses in a binary decision.

As shown in Fig. 1, the neural architecture starts with a convolution layer, which uses a kernel size of 64 ×9 and a stride of 64 × 1. The convolution layer has a rectifying linear unit (ReLU) activation function, and is followed by an average pooling layer with a 1 × 256 kernel, and two FC layers with 20 and 10 neurons, respectively. Finally, a softmax output layer is added for binary decision.

During training, we adopt the cross-entropy loss function as the cost function in the adaptive moment estimation algorithm (Adam) [20]. The learning rate is set to 1×10^*-*3^. The channel attention mechanism and the CNN classifier are jointly trained as a single system. At run-time, the system produces two values at the output nodes for decision making.

## III. Experiments and Results

### A. Experimental setup

In this study, experiments were carried out on a public auditory attention detection dataset [21], recorded at KU Leuven, which is referred to as KUL Dataset. Briefly, 64-channel EEG data was recorded from 8 male and 8 female normal-hearing subjects when they listened to two competing speakers, and were instructed to attend to one speaker. EEG data was recorded at a sample rate of 8192 Hz using a BioSemi ActiveTwo system. Four Dutch short stories, narrated by different male speakers, were used as speech stimuli through a pair of insert earphones. The whole experiment was split into 8 trials and each trial lasts 6 minutes. The auditory stimuli were presented from 90° to the left and 90° to the right of the subject, respectively. Overall, the EEG data from 16 normal-hearing subjects was collected, and there was 48 minutes of data for each subject.

### B. Data processing

The EEG data of each channel were firstly re-referenced to the mean of the response of all channels. Then, all the EEG data were bandpass filtered between 1 and 50 Hz, and subsequently down-sampled to 128 Hz. The speech stimuli were firstly passed through a Gammatone filterbank ranging from 150 Hz to 8 kHz. All of the sub-bands were power-law compression with 0.6 [22]. Finally, the speech envelopes were transformed into their respective absolute envelopes by a Hilbert transformation, low-pass filtered with 50 Hz and down-sampled from 512 Hz to 128 Hz to match the EEG data.

The data set was randomly split into a training set (60%) and a validation set (20%), and a test set (20%). For each partition, data segments were generated with a sliding window, denoted as the decision window, with a overlap of 50%. We maintain a balanced number of speaker A/B attention samples by subject, i.e, a random guess will give a 50% accuracy. All the repetitions were discarded to keep the training, validation, and test set mutually exclusive. We are particularly interested in low-latency attention detection, therefore, we only report the detection accuracy for four short decision windows: 0.1, 0.5, 1.0, and 2.0 second. To avoid initialization bias, the experiments of each subject were carried out 10 times with random initialization to report a subject average accuracy.

### C. Experiment results

A comprehensive comparative study was carried out. We re-implemented two reference baselines, namely the stimulus reconstruction (linear) model [5], and the CNN (non-linear) model [10], on KUL Dataset with 64-channel EEG. The difference between our CNN-CM model and the CNN baseline lies in the additional channel attention mechanism.

As shown in Table I, the CNN-CM model significantly outperforms the linear decoder with a large margin for all decision windows. These results corroborate with previous studies [7], [10], [11], [12]. It is noted that the detection accuracy increases as we increase the decision window size. Encouragingly, the CNN-CM model has seen a mean accuracy of 77.2% (SD: 8.24) for 0.1-second decision window, which represents an improvement of 15.9% over the linear baseline for 2-second decision window. Moreover, the average detection accuracy of the CNN-CM model exceeded 86% at a temporal resolution of around 1 second, comparable to the human’s time lag when switching attention [23]. We are not aware of other decoders that achieve similar accuracy under such low latency settings.

**TABLE I.**
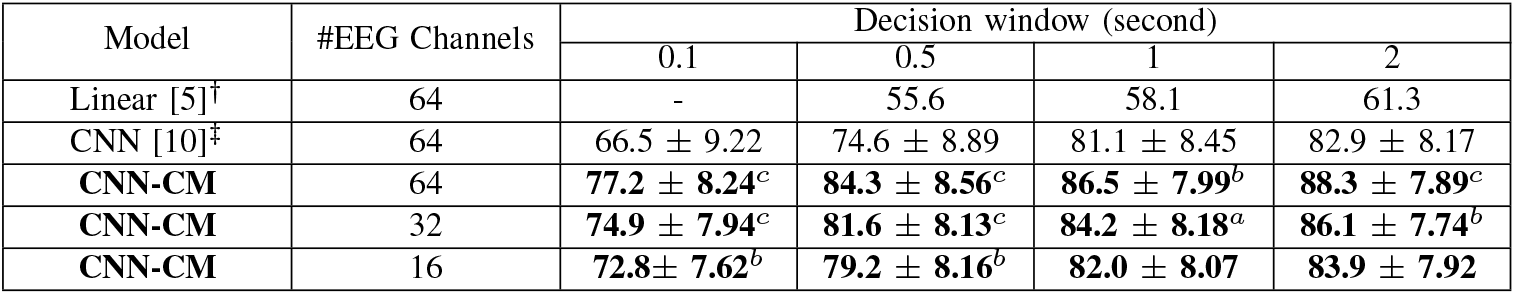
Auditory attention detection accuracy (%) and its standard deviation (*±*) in a comparative study. † denotes our re-implementation of the linear model in [5]. ‡ denotes our re-implementation of the CNN model in [10]. ^*a, b, c*^ denote the significant increase of AAD accuracy over the CNN method [10] with *p* <0.05, *p* <0.01, and *p* <0.001 respectively.

As shown in Fig. 2, the CNN-CM model outperforms the non-linear CNN model in [10] with consistent improvements in AAD accuracy with 5.4% for 1.0 and 2.0 second decision windows, 9.7% for 0.5-second decision window, and 10.7% for 0.1-second decision window, respectively. With the channel attention mechanism, our CNN-CM model significantly outperforms the CNN model (paired *t*-test: *p* <0.01).

**Fig. 2.**
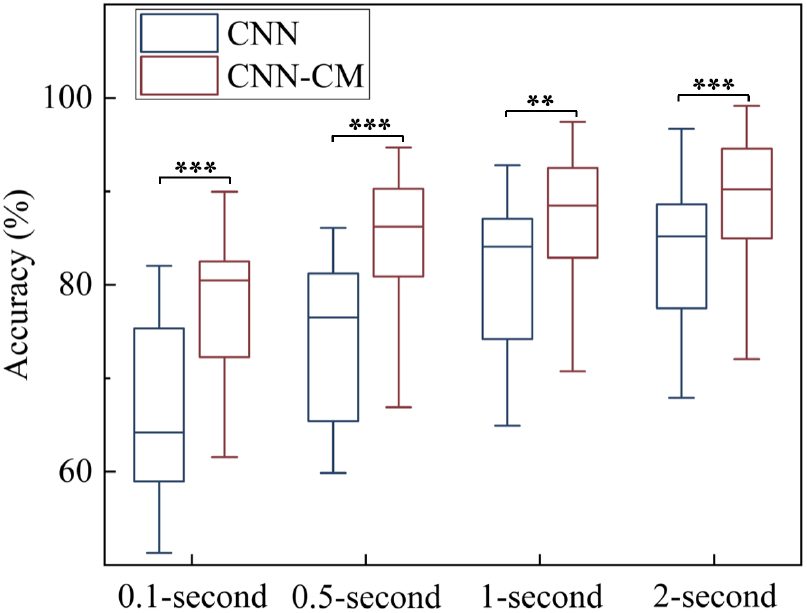
Auditory attention detection accuracy (%) of CNN-CM and CNN models with 64-channel EEG for different decision windows. Statistically significant differences: \*\**p* <0.01, \*\*\**p* <0.001. It is observed that a larger decision window leads to a higher accuracy with a lower variance.

### D. Effect of electrode reduction

We would like to examine if the proposed channel attention mechanism is still effective with a reduced number of EEG signals.

Specifically, we reduced the EEG signals from 64-channel to 32-channel and 16-channel following the electrode locations of the international 10/20 system, respectively. As summarized in Table I, though AAD performance decreases with low-density EEG systems [13], the CNN-CM model continues to outperform both linear and non-linear baselines. It is noteworthy that the CNN-CM model with 16-channel EEG achieves even better results than the state-of-the-art CNN model across all different decision windows.

Considering that a low number of EEG electrodes can considerably reduces preparation time and help to meet the requirements for BCI-based applications, our CNN-CM model are more suitable for neuro-steered hearing prostheses.

## IV. Discussion and Conclusions

All experiments have confirmed the effectiveness of the proposed channel attention mechanism. We consider that two main factors have contributed to the significant improvement of the CNN-CM model over the baselines. One is the task-oriented feature representation with channel attention mask. The attention mechanism dynamically focuses the attention on effective EEG channels, as evidenced by the reduced standard deviation of accuracy across the participating subjects over the CNN baseline in Table I. Another is the joint training between the channel attention mechanism and the CNN classifier, that allows for feature representation and classifier to be optimized for attention detection performance.

## ACKNOWLEDGMENT

This research work is supported by Programmatic Grant No. A18A2b0046 and A1687b0033 from the Singapore Government’s Research, Innovation and Enterprise 2020 plan (Advanced Manufacturing and Engineering domain).

The work by Haizhou Li is also funded by the Deutsche Forschungsgemeinschaft (DFG, German Research Foundation) under Germany’s Excellence Strategy (University Allowance, EXC 2077, University of Bremen, Germany).

The work by Longhan Xie is also funded by the National Natural Science Foundation of China (Grant No. 52075177).

## Notes

### Competing Interest Statement

The authors have declared no competing interest.

